# Offline Two-dimensional Liquid Chromatography-Mass Spectrometry for Deep Annotation of the Fecal Metabolome

**DOI:** 10.1101/2023.05.31.543178

**Authors:** Brady G. Anderson, Alexander Raskind, Rylan Hissong, Michael K. Dougherty, Sarah K. McGill, Ajay Gulati, Casey M. Theriot, Robert T. Kennedy, Charles R. Evans

## Abstract

Compound identification is an essential task in the workflow of untargeted metabolomics since the interpretation of the data in a biological context depends on the correct assignment of chemical identities to the features it contains. Current techniques fall short of identifying all or even most observable features in untargeted metabolomics data, even after rigorous data cleaning approaches to remove degenerate features are applied. Hence, new strategies are required to annotate the metabolome more deeply and accurately. The human fecal metabolome, which is the focus of substantial biomedical interest, is a more complex, more variable, yet lesser-investigated sample matrix compared to widely studied sample types like human plasma. This manuscript describes a novel experimental strategy using multidimensional chromatography to facilitate compound identification in untargeted metabolomics. Pooled fecal metabolite extract samples were fractionated using offline semi-preparative liquid chromatography. The resulting fractions were analyzed by an orthogonal LC-MS/MS method, and the data were searched against commercial, public, and local spectral libraries. Multidimensional chromatography yielded more than a 3-fold improvement in identified compounds compared to the typical single-dimensional LC-MS/MS approach and successfully identified several rare and novel compounds, including atypical conjugated bile acid species. Most features identified by the new approach could be matched to features that were detectable but not identifiable in the original single-dimension LC-MS data. Overall, our approach represents a powerful strategy for deeper annotation of the metabolome that can be implemented with commercially-available instrumentation, and should apply to any dataset requiring deeper annotation of the metabolome.

## INTRODUCTION

Untargeted metabolomics data typically contains hundreds to thousands of unknown features, even after rigorous data reduction techniques are applied. The most important and readily available method to help identify biologically relevant unknown metabolites is the acquisition of tandem mass spectrometry (MS/MS) data, which can be searched against experimental and *in-silico* spectral databases.^5^ Yet even with an assortment of spectral libraries containing thousands of high-quality spectra, some features are inherently difficult to identify because the MS/MS spectra they produce contain few unique product ions by collision-induced dissociation.^6,7^ Separation before MS detection by liquid chromatography (LC) can alleviate ionization competition and elevate MS signal, yet many minor features and corresponding fragment ions remain too low in abundance to enable confident identification by data-dependent or data-independent MS/MS acquisition.^8^

To adequately sample polar and nonpolar metabolites, untargeted metabolomics studies commonly require multiple separation methods, including hydrophilic interaction liquid chromatography (HILIC) and reversed-phase LC (RPLC). Therefore, experiments with numerous samples must be performed using relatively short run times (∼20 min or less) to achieve adequate throughput. To achieve accurate quantification, they must also use sample loading quantities that avoid column or detector saturation for the most abundant features. However, these conditions are not optimized for compound identification. Previously, our group demonstrated a nearly 10-fold improvement in identification could be achieved when longer run times (∼3 hours), higher sample loading, and multi-run precursor ion exclusion were used.^9^ Such conditions only need to be used for the analysis of a single representative pooled sample since the software tool *metabCombiner* allowed confident alignment of newly-identified compounds with features detectable but not identifiable under conventional LC-MS conditions.^10^ However, at the longest gradients and highest loading conditions, broader peaks and increased coelution resulted in a leveling-off of the number of compounds identified in a run, suggesting that a practical limit exists for one-dimensional (1d) separations on analytical bore columns (∼2.1 mm i.d.).

Larger diameter columns offer higher loading capacity, but since ESI sensitivity is mostly concentration-dependent, higher mass loading can only be translated to improved sensitivity when eluted compounds can be concentrated before re-injection on the MS.^11^ By performing a separation using a semi-preparatory (10 mm i.d.) column, collecting fractions, then concentrating the fractions and re-injecting on an analytical column, higher mass loading can be achieved without overloading the analytical column to the extent that would occur if the same amount of unfractionated sample had been injected on the analytical column. Fractionating a sample before a second-dimension separation further reduces coelution and corresponding ion suppression.^12^ One approach utilizing a semi-preparatory ion-pairing RPLC first dimension separation detected 3564 unique ion pairs in human urine after collected fractions were isotopically dansylated to impart additional hydrophobicity prior to an analytical scale RPLC separation.^13^ In a 1d RPLC-MS separation of the same sample, only 1218 ion pairs were observed. A similar approach was recently published utilizing supercritical fluid chromatography (SFC) for the first-dimension separation of lipid classes.^14^ SFC as a first-dimension separation is potentially advantageous over semi-preparatory LC as faster flow rates, shorter fraction drying times, and more efficient separations can be achieved, although it is less suitable for analyzing water-soluble metabolites.^7,15^ In this approach, 404 lipids were identified with the SFC fractionation workflow compared to 150 with a 1d RPLC-MS approach. Various column chemistry combinations have also been evaluated for orthogonality and application to compound identification.^16,17^ RPLC x HILIC and ion exchange (IEX) x RPLC configurations yielded the greatest identification performance for urine metabolites.^16^ Taken together, these studies provide evidence for the potential of offline two-dimensional chromatography to improve the detection and identification of low-abundance metabolites.

Most, though not all, sample types commonly investigated in metabolomics are amenable to the moderate degree of scale-up required to gain sensitivity using offline LC x LC. Among suitable sample types, human plasma and urine have been extensively evaluated using a multitude of techniques, including multidimensional chromatography.^9,18–21^ In contrast, the fecal metabolome remains substantially less well characterized, even as interest grows in studying the gut microbiome and its interaction with the host’s metabolism.^22,23^ This study demonstrates an offline two-dimensional (LCxLC) MS/MS metabolomics approach to identify low abundance human fecal metabolites. Multiple criteria were used to assess the accuracy and confidence of these compound identifications, including MS/MS database search score thresholds and RT alignment with either experimentally analyzed authentic standards or computationally predicted RT for novel compounds. In comparison to conventional (20-minute) RPLC and HILIC separations, the offline two-dimensional methods (RPLC x HILIC and RPLC x RPLC) more than doubled the number of unique database match assignments (1513 to 3414). Our study demonstrates substantial improvement in metabolite identification can be achieved using fractionation and preconcentration of complex samples prior to untargeted metabolomics analysis.

## EXPERIMENTAL SECTION

### Materials

LC-MS grade acetone, acetonitrile, methanol, and water were purchased from Fisher Scientific. LC-MS grade formic acid, ammonium formate, ammonium acetate, and 25% by weight ammonium hydroxide mobile phase modifiers were purchased from Sigma-Aldrich. Internal standards were acquired from Cambridge Isotope Laboratories (Andover, MA). Human metabolite standards were purchased from MetaSci (Richmond Hill, Ontario). Fecal samples were pooled from multiple de-identified human subjects collected under informed consent under IRB #16-2283 at the University of North Carolina Hospital. Fecal samples were aliquoted into Eppendorf tubes and stored at −80 °C until extraction.

### Sample Extraction

Fecal samples were weighed into pre-tared 2 mL Precellys® (Bertin Corp.) compatible vials, and one 2.8 mm stainless steel bead was added to aid homogenization. Fecal matter was homogenized using a Precellys® Evolution by two 20 s cycles separated by a 30 s break. Extraction solvent was used at a ratio of 1 mL per 5 g feces and was comprised of 1:1:1 methanol:acetonitrile:acetone containing 10 μM of D_3_-creatine, D_10_-isoleucine, D_2_-biotin, D_5_-tryptophan, D_3_-caffeine, D_3_-octanoylcarnitine, D_3_-palmitoylcarnitine, D_4_-deoxycholic acid, D_4_-cholic acid, and D_7_-arginine as internal standards. Following extraction, samples were centrifuged for 10 min at 17,000 rpm. 100 μL aliquots of supernatant were transferred to Eppendorf vials, dried under a gentle stream of nitrogen, and stored at −80 °C. On the day of analysis, the dried extracts were reconstituted in 85:15 acetonitrile:water for HILIC analysis or 9:1 water:methanol for RPLC analysis with volumes as described below. Pooled samples were prepared by combining equal volumes of reconstituted fecal matter extracts from all subjects.

### Sample Fractionation

For pooled samples analyzed using offline two-dimensional chromatography, semi-preparative RPLC separations of 2-fold concentrated fecal matter extract (1800 μL total dried extract reconstituted in 900 μL 9:1 water: methanol) were performed on a Waters Atlantis T3 C18 column (10 x 150 mm; 5 μm). The column compartment was maintained at 55 °C. Mobile phases consisted of water with 0.1% v/v formic acid and methanol with 0.025% v/v formic acid and utilized the following gradient: 0-1 min 0% B; 1-20 min 100% B; 20-40 min 100% B. The flow rate was set to 3 mL/min and LC effluent was split 50:1 between the fraction collector and an Agilent 6520 quadrupole time of flight mass (QTOF) spectrometer. The MS parameters for the QTOF were set as the following: Ionization mode, positive; Mass range, 100-1700 m/z; Gas temperature, 325 °C; Drying gas, 5 L/min; Nebulizer, 30 psig; Capillary voltage, 3500 V; Fragmentor 175 V; Skimmer, 65 V; Octupole 1 RF Vpp, 750 V; Acquisition rate, 1.37 spectra/s; Acquisition time, 1000 ms/spectrum; Spectrum data type, centroid.

Ninety-four 0.35-minute fractions with a volume of 1.05 mL each were collected into tapered-base glass autosampler vials (Thermo Scientific) beginning at 3.0 minutes post-injection of the semi-preparative method. The fractions were then dried using a GeneVac® Ez-2 (Ipswich, United Kingdom) vacuum centrifuge. The dried fractions were reconstituted in 50 μL of method-specific reconstitution solvent (RPLC/HILIC). Adjacent fractions were combined, resulting in forty-seven 100 μL fractions. 5 μL of each fraction was subsequently analyzed by analytical LC-MS/MS methods described below.

### Analytical Separations

Individual subject samples, fractionated and unfractionated pooled samples, and analytical standards were analyzed by HILIC (Waters BEH Amide, 2.1 x 100 mm, 1.7 μm) and RPLC at high pH (Waters Charged-Surface Hybrid [CSH] C18, 2.1 x 100 mm, 1.7 μm) in both positive and negative ion modes on a Thermo Vanquish Horizon LC coupled to an Orbitrap ID-X mass spectrometer. For HILIC separations, mobile phase A consisted of 95:5 water:acetonitrile with 10 mM ammonium formate plus 0.125 % v/v formic acid and mobile phase B was 5:95 water:acetonitrile with the same additive concentrations. HILIC separations utilized the following gradient: 0 min, 100% B; 0-0.5 min 100% B; 0.5-7 min 85% B; 7-9 min 85% B; 9-16 min 50% B; 16-16.1 min 100% B; 16.1-20 min 100% B. For CSH separations, mobile phase A consisted of water with 10 mM ammonium acetate plus 0.025% ammonium hydroxide (v/v) and mobile phase B was methanol with the same additives. CSH separations utilized the following gradient: 0 min 0% B, 0-5 min 60% B; 5-13 min 99% B; 13-17 min 99% B; 17-17.1 min 0% B; 17-20 min 0% B. Mass spectrometer settings for both ionization modes were as follows: sheath gas, 40; aux gas, 10; sweep gas, 1; ion transfer tube temp, 325 °C; vaporizer temp, 300 °C; orbitrap resolution. 120000; scan range, 70-800 m/z; RF lens, 45%; normalized AGC target, 25%; maximum injection time, auto; microscans, 1; data type, profile; internal mass calibration, EASY-IC^TM^. Positive ion spray voltage was set to 3200 V while negative ion mode was set to −3200 V. Instrument settings for MS^1^ and MS/MS methods were identical, except orbitrap resolution was decreased to maximize MS/MS spectra collection. The data-dependent MS/MS methods utilized the following settings: full scan orbitrap resolution, 60000; intensity threshold, 1.0×10^4^; dynamic exclusion properties; exclusion duration 3 seconds (exclude after 1 time with ± 5 ppm); isolation mode, quadrupole; isolation window, 1.2 m/z; activation type, HCD; collision energy mode, assisted; collision energies, 20, 40, and 80%; detector type, orbitrap; orbitrap resolution, 30000; normalized AGC target, 20%; maximum injection time, 54 ms; microscans, 1; data type, centroid; cycle time, 1.2 s.

### Data Processing and Compound Identification

MS^1^ feature detection and adduct annotation for individual samples was performed in Thermo Compound Discoverer 3.3 with the following settings: minimum peak count, 1; mass tolerance, 5 ppm; minimum peak intensity, 10000; minimum number of scans per peak, 5; S/N threshold, 1.5; gap ratio threshold, 0.35; peak width, 0.1-1.5 min; grouping tolerance, 5 ppm and 0.3 min. An exported feature table with peak areas was then formatted for analysis with MetaboAnalyst 5.0.^24^ A statistical one-factor analysis of the unpaired samples was performed. Sample data was normalized by the median, logarithmically transformed (base-10), and statistically relevant features were classified as a minimum of ± 1.25-fold change and FDR-corrected threshold of 0.1 or lower. Peak areas for the internal standards were checked before and after normalization.

Our software tool *MetIDTracker* was utilized for MS/MS database searching, manual spectral review when needed, and alignment of features of interest from the MS^1^ data.^9^ Libraries used for spectral search were NIST20 (www.nist.gov), MassBank of North America (MONA, massbank.us), and the MS-Dial fork of LipidBlast.^25^ “Identity,” “in-source,” and “hybrid” searching strategies were used sequentially to match spectra to a specific structure when possible and otherwise to annotate features by compound class as described previously.^9^ A spectral entropy scoring algorithm developed by Li et al.^26^ has additionally been incorporated into *MetIDTracker;* this method has been demonstrated to produce a *false discovery rate* under 10% (one-bond isomers are not penalized). Other MS/MS metrics, including NIST score, dot product, reverse dot product, total intensity, and pattern recognition entropy, are also calculated. As an additional metric of spectral quality, precursor ion purity is calculated as the ratio of the precursor ion intensity over the summed intensity of all ions within the quadrupole isolation window in the MS^1^ scan.

Standards from the MetaSci human metabolite library (∼1000 endogenous and human exposome metabolites) were multiplexed into mixtures containing 50 or fewer non-isobaric compounds and retention times (RT) were determined for all observed standards in both HILIC and CSH modes using Skyline.^27^ Retention time prediction (RTP) models were trained and validated in R package Retip with an 80:20 random training:testing split.^28^ The extreme gradient boosting RTP model for the HILIC and CSH analyses with the lowest root mean square error from five trials was used to predict retention times of proposed database matches.

Using RTP in addition to MS/MS database matches can improve confidence in the validity of a compound identification, though not to the same extent as matching the experimentally measured retention time of an authentic standard. Therefore, only features satisfying MS/MS search score criteria (entropy score greater than or equal to 0.65) and with close RT alignment (± 0.5 min) to the matching analytical standard were treated as MSI level 1 (MSI1) identifications. Similarly, MS/MS “hits” that did not align with an available standard were downgraded to compound class-level annotations (MSI3). Spectral hits for compounds not matching an analytical standard but with a measured retention time within ± 1.0 min of retention time predicted by Retip were classified as “MSI2A” identifications. Features matched as identity hits by MS/MS entropy score alone were termed MSI2B IDs. MSI3 annotations were assigned for features meeting one or more of the following criteria: NIST hybrid score ≥ 600, 0.5 ≤ identity MS/MS entropy score < 0.65, or in-source MS/MS entropy score ≥ 0.65.^29^ Remaining detected features were classified as MSI level 4 (unknowns). MSI3 and MSI4 features were further consolidated to the highest intensity MS/MS feature within 0.2 minutes and ± 0.0025 m/z. The number of identifications by each method was counted by the number of unique first 14 characters of an InChIKey with an MSI1, MSI2A, or MSI2B identification level. Proposed identifications of the same compound that existed in four or more fractions were considered background ions and removed from consideration.

## RESULTS AND DISCUSSION

### Evaluating Samples with Conventional LC-MS Metabolomics

Prior to fractionation, individual subject samples used to generate the pooled sample were analyzed by CSH and HILIC LC-MS methods as described above. Total ion chromatograms (TIC) of the fecal extracts under different modes of chromatography and ionization are shown in Figure 1. High variability in the inter-subject fecal metabolome is expected due to differences in diet, microbiome, and other factors.^32,33^ One notable aspect was the presence of sizable clusters of peaks in the CSH separation attributed to polyethylene glycol (PEG). These were likely present due to osmotic laxative intake by some subjects.^34,35^ PEG-related ions were recorded and ignored in subsequent analysis, although ion suppression of co-eluting metabolites may have altered the apparent abundance of differential metabolites in this retention time range.^36^

**Figure 1.**
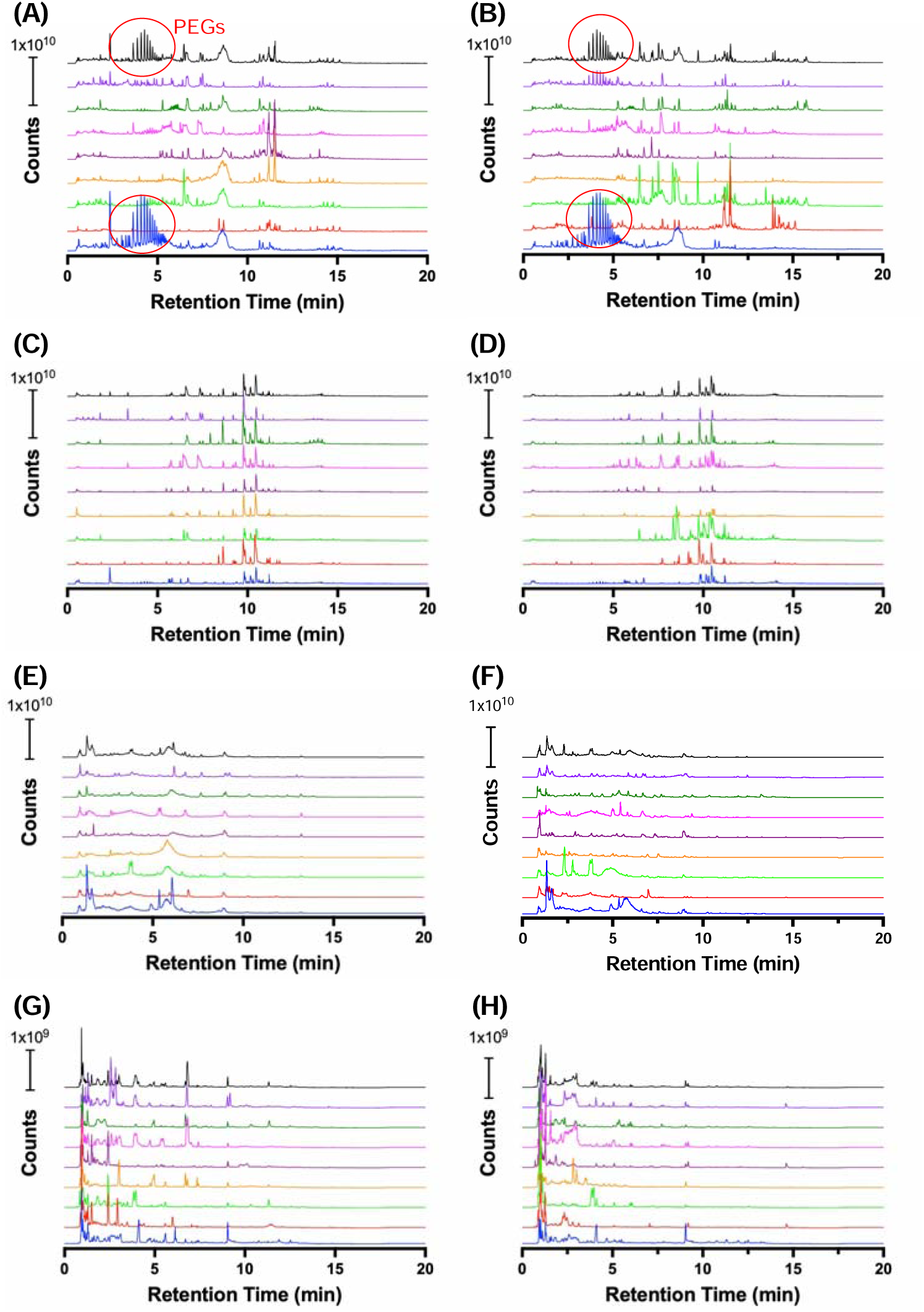
TICs of individual subject samples. Chromatograms were collected by (A-B) CSH positive mode, (C-D) CSH negative mode, (E-F) HILIC positive mode, and (G-H) HILIC negative mode.

### Assigning Confidence to Metabolite Identifications

Several classification schemes have been proposed to report identification confidence for metabolomics data, however, assignment to a specific level in any such scheme is subject to interpretation. For instance, an MSI2 identification typically requires a MS/MS spectral match, but no specific search algorithm or score threshold is specified.^37–39^ Higher confidence identifications can be achieved when multiple pieces of experimental evidence including retention time alignment are used. Many laboratories have developed extensive retention time libraries based on authentic standards for this reason. However, the financial resources required to establish these libraries can be significant, and data acquisition method details are not always transparent. To assess identification confidence to metabolites not included in the library, retention time prediction (RTP) CSH and HILIC models were created with R package Retip.^28^ Replicate RTP model performance is reported in Tables 1-2 and Figure 2. Modeling HILIC retention was less accurate than CSH retention likely due to a more complex mechanism of separation.^40–42^ It is our opinion that RTP accuracy is not sufficient to confidently confirm identifications, but it adds supporting evidence if observed and predicted retention times align within an empirically-determined margin, which was selected as ± 1 min for this study.

**Figure 2.**
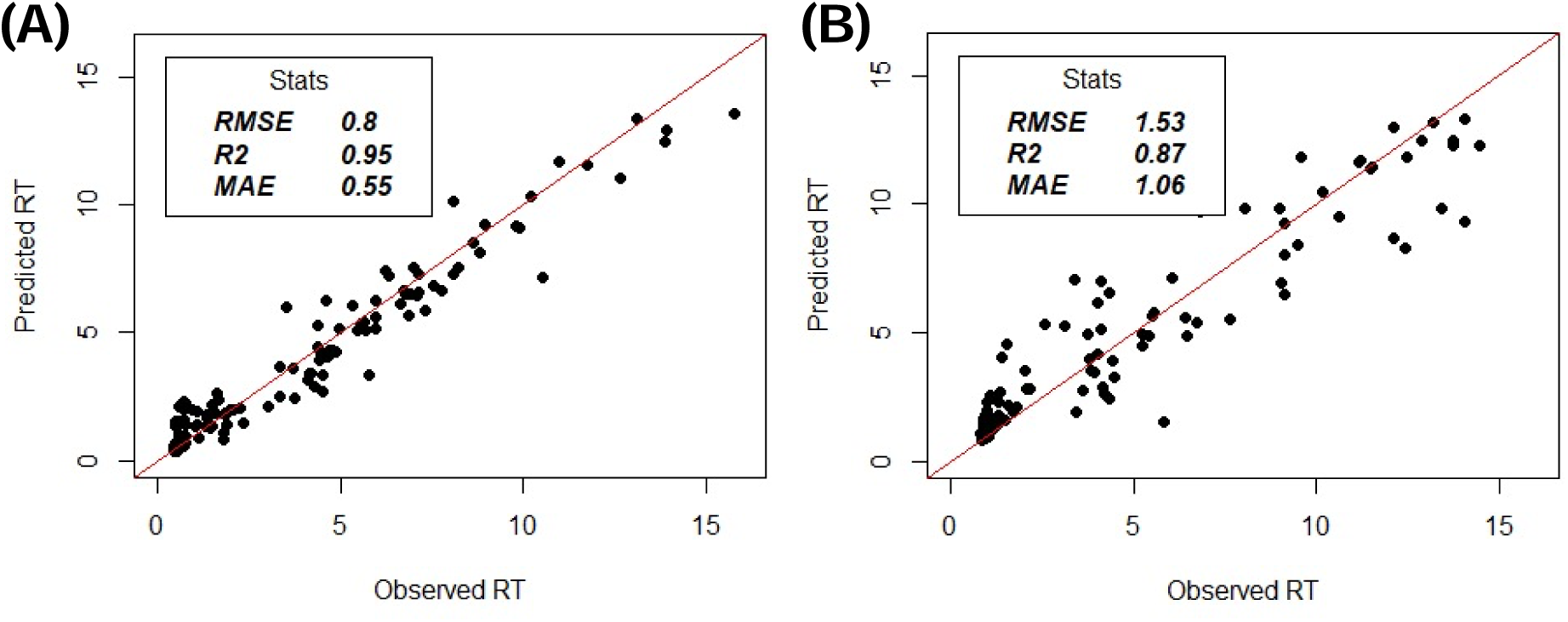
Observed versus predicted retention time for the best (A) CSH and (B) retention time models. Root mean square error (RMSE), linear regression correlation (R2), and mean average error (MAE) are shown for each model.

**Table 1.** Retention time prediction model replicates for the CSH method. The model highlighted in red had the lowest root mean square error and was utilized for predicting retention times.

| Model Name | Root mean square error (min) | R2 (observed vs predicted RT) | Mean average error (min) | 95 % confidence interval (+/- min) |
| --- | --- | --- | --- | --- |
| **Xgb** | **0.8** | **0.95** | **0.55** | **1.38** |
| Xgb2 | 1.05 | 0.91 | 0.62 | 1.79 |
| Xgb3 | 0.82 | 0.94 | 0.55 | 1.39 |
| Xgb4 | 0.94 | 0.90 | 0.58 | 1.51 |
| Xgb5 | 0.82 | 0.93 | 0.58 | 1.22 |

**Table 2.** Retention time prediction model replicates for the HILIC method. The model highlighted in red had the lowest root mean square error and was utilized for predicting retention times.

| Model Name | Root mean square error (min) | R2 (observed vs predicted RT) | Mean average error (min) | 95 % confidence interval (+/- min) |
| --- | --- | --- | --- | --- |
| Xgb | 1.94 | 0.76 | 1.27 | 2.75 |
| **Xgb2** | **1.53** | **0.87** | **1.06** | **2.41** |
| Xgb3 | 1.88 | 0.84 | 1.24 | 3.07 |
| Xgb4 | 2.16 | 0.76 | 1.26 | 3.54 |
| Xgb5 | 1.76 | 0.86 | 1.21 | 2.96 |

The lack of validated methods to estimate false discovery rates for compound identification in metabolomics leaves manual review as the only practical option to assess hit quality; however, it is inherently subjective. A new spectral entropy scoring algorithm has been demonstrated to lower false positive rates.^26^ However, with the spectral entropy thresholds suggested in the literature (0.75), we observed many metabolite identifications with retention time alignment to authentic standards or predicted retention times that otherwise would not have been confirmed. We therefore selected an entropy score threshold of 0.65 as a good balance between identification confidence and false discovery. An example head-to-tail plot of diltiazem with an entropy score of 0.681 and retention time alignment to a pure standard is shown in Figure 3.

**Figure 3.**
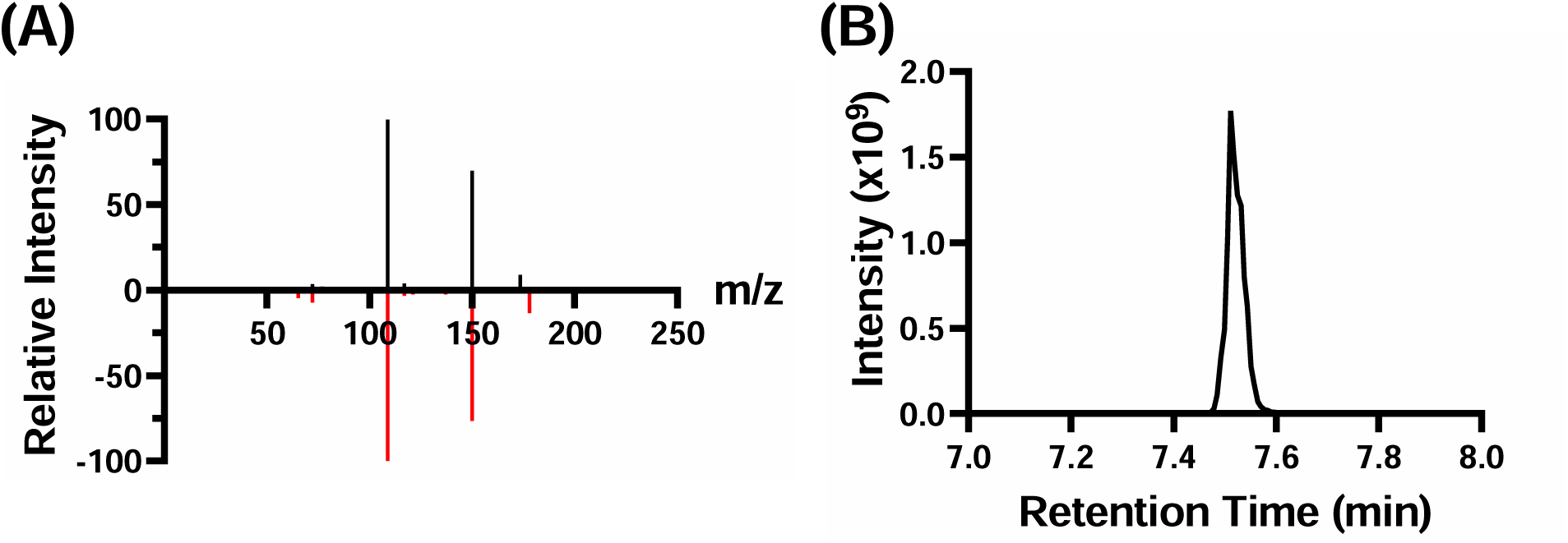
(A) Head-to-tail plot of and (B) extracted ion chromatogram of diltiazem ([M+H]^+^). Experimental and library spectral alignment had an entropy score of 0.681.

### Assessing Identification Potential for 1d and 2d Approaches

To properly evaluate the potential of offline 2d separations and justify the increased time and sample consumption they require, identification performance must be assessed relative to typical 1d separations. An exclusion list of non-relevant precursor ions was first created by running a CSH and HILIC blank which consisted of the corresponding reconstitution solvents. Then five iterative MS/MS injections with run-to-run precursor ion exclusion enabled were performed using a pooled fecal extract, and compounds identified according to criteria described in the methods were tabulated (Figure 4). The number of unique identifications increased with each successive injection but trended toward a plateau for both ionization modes and separations. 931 and 288 total identifications (MSI1 through MSI2B) were observed for the CSH separations in positive and negative mode, and 545 and 292 were observed using HILIC. Lower identification totals for negative mode were expected, as the acidity of the mobile phases favored the formation of positive ions; this compromise was accepted to allow retention time alignment between ionization modes. 1147 unique identifications were made by CSH and 770 by HILIC when positive and negative mode results were combined; 1513 unique identifications were made in total for all 1d methods. MSI-level feature breakdowns for each 1d method are illustrated in Figure 5. Greater than 65% of collected MS/MS features for all methods were classified as unknowns.

**Figure 4.**
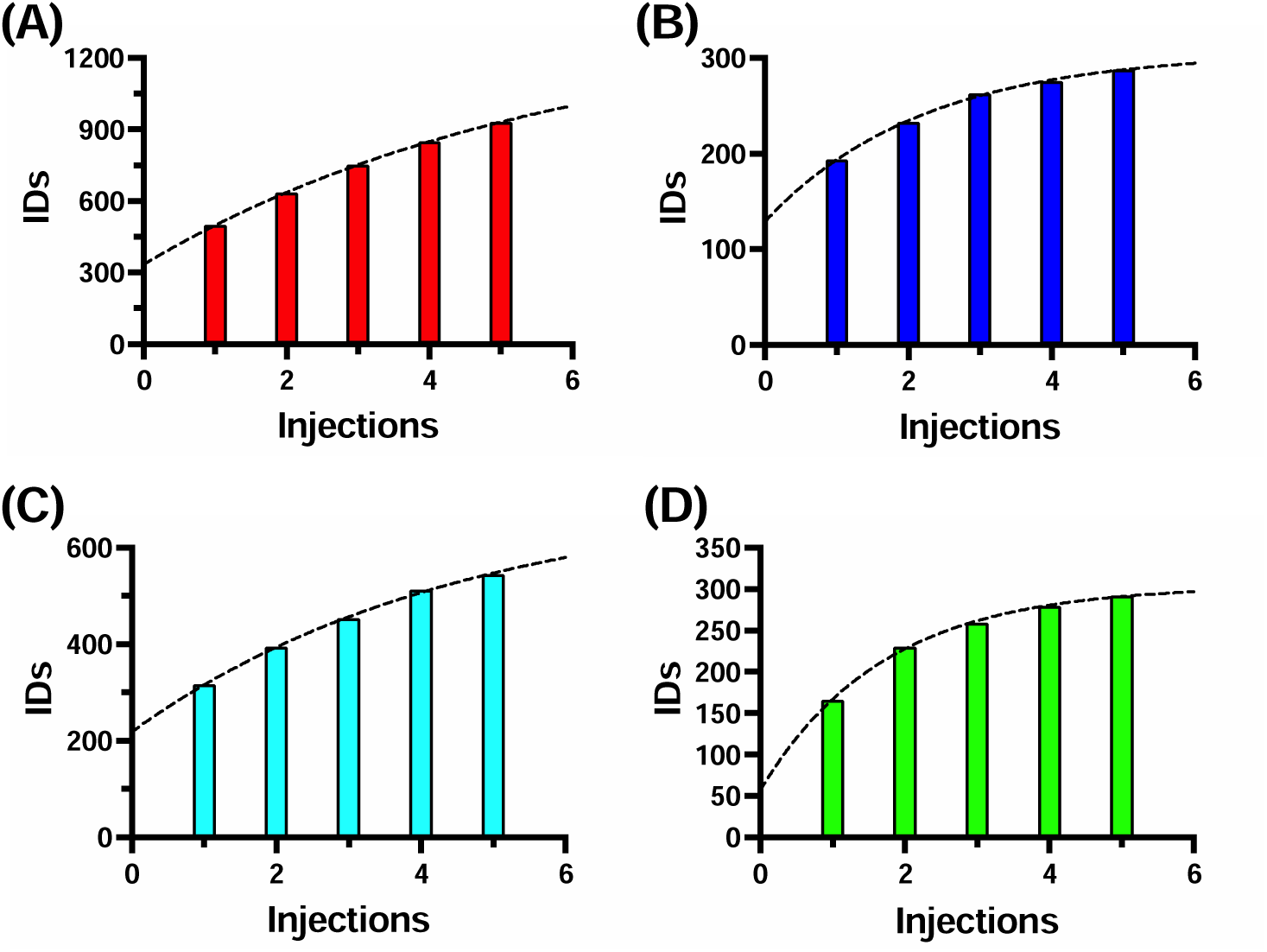
The number of cumulative unique identifications made by five iterative injections of (A) CSH positive mode, (B) CSH negative mode, (C) HILIC positive mode, and (D) HILIC negative mode methods.

**Figure 5.**
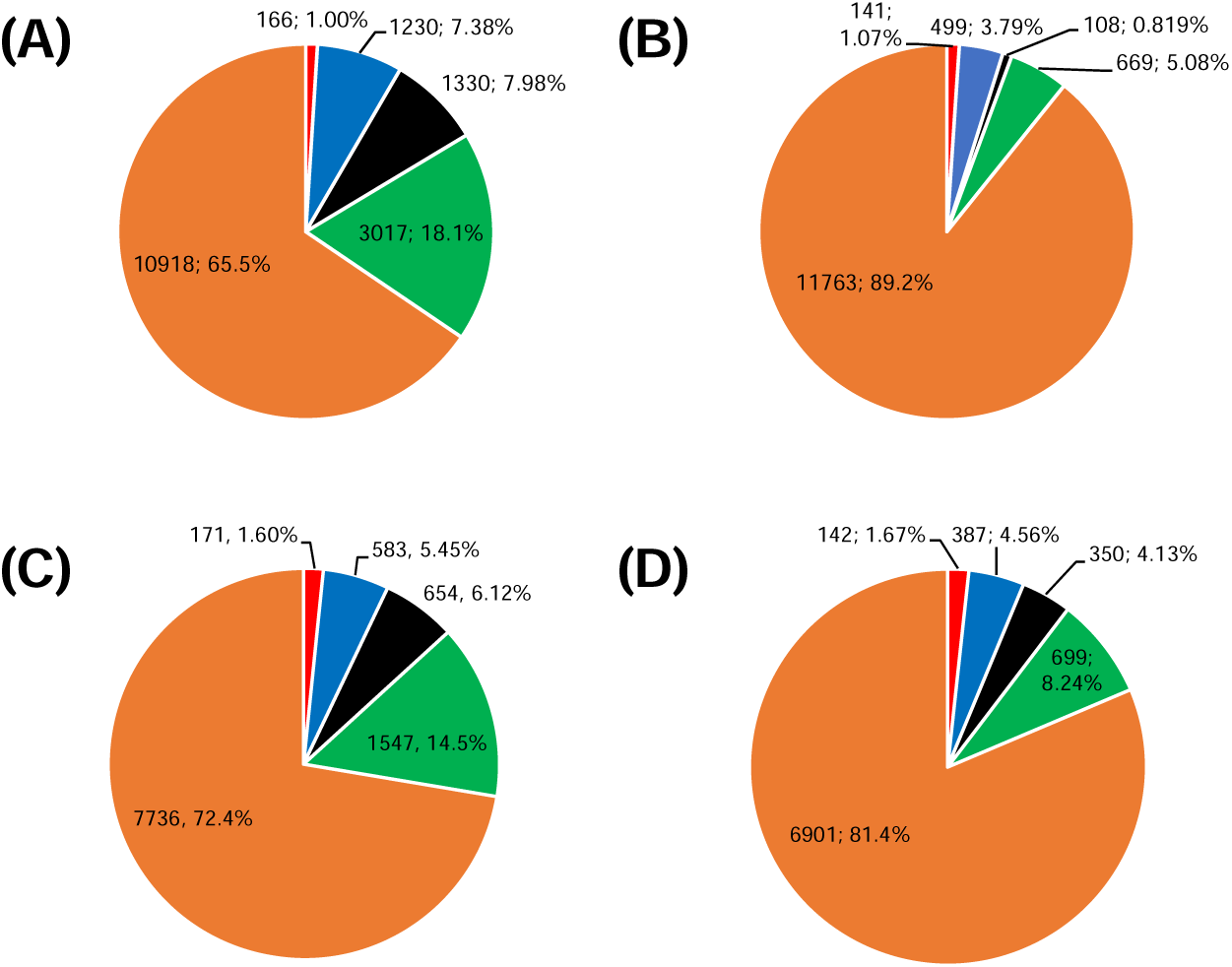
MSI level feature consolidation for (A) CSH positive mode, (B) CSH negative mode, (C) HILIC positive mode, and (D) HILIC negative mode methods. The number of MSMS features and overall percentage are displayed. ID levels MSI1, MSI2A, MSI2B, MSI3, and MSI4 are shown in red, blue, black, green, and orange.

A low pH semi-preparative RPLC first dimension separation was used for all offline 2d analyses as RPLC supports higher loading capacity than HILIC, making HILIC a less ideal choice to produce high-concentration fractions for downstream analysis.^43^ The effective peak capacity of offline two-dimensional (2d) separations is determined by several factors, including the peak capacity of each separation, separation method orthogonality, and fraction sampling frequency.^44,45^ A high-pH RPLC second dimension separation (CSH) was thus employed as the second dimension to generate orthogonality in our RPLC x RPLC method; however, RPLC x HILIC separations are generally more orthogonal and higher total peak capacities can be realized. Several previous studies have systematically investigated column combinations and orthogonality in greater detail;^16,17^ we report here the potential for enhanced metabolite identification and unidentified compound classification using selected chromatographic methods after first simplifying an extract by fractionation and preconcentration.

A TIC from the first-dimension semi-preparative RPLC separation is shown in Figure 6, while second-dimension TICs of the fractions are displayed as heat maps with intensity as a logarithmically scaled grayscale gradient in Figure 7. A clear diagonal band of peaks can be seen for the CSH second-dimension separations, signifying low orthogonality with the first-dimension separation. For HILIC second-dimension separations, earlier eluting (polar) compounds from the RPLC first dimension were, as expected, spread over more of the total separation space, whereas late-eluting (nonpolar) compounds were lightly retained and eluted early in the HILIC gradient. The number of unique identifications made per fraction (Figure 8) further shows this discrepancy in orthogonality, as more identifications for the first ∼20 fractions were observed with the RPLC x HILIC combination. In total, 1479, 555, 1917, and 830 identifications were made with the fractionation approach by a CSH positive mode, CSH negative mode, HILIC positive mode, and HILIC negative mode. 1917 and 2548 unique identifications in the fractions were made in total for both modes of the CSH and HILIC methods, and 3414 total unique identifications were achieved across all methods. Figure 9 summarizes the total number of identifications made by each of the 1d and 2d methods.

**Figure 6.**
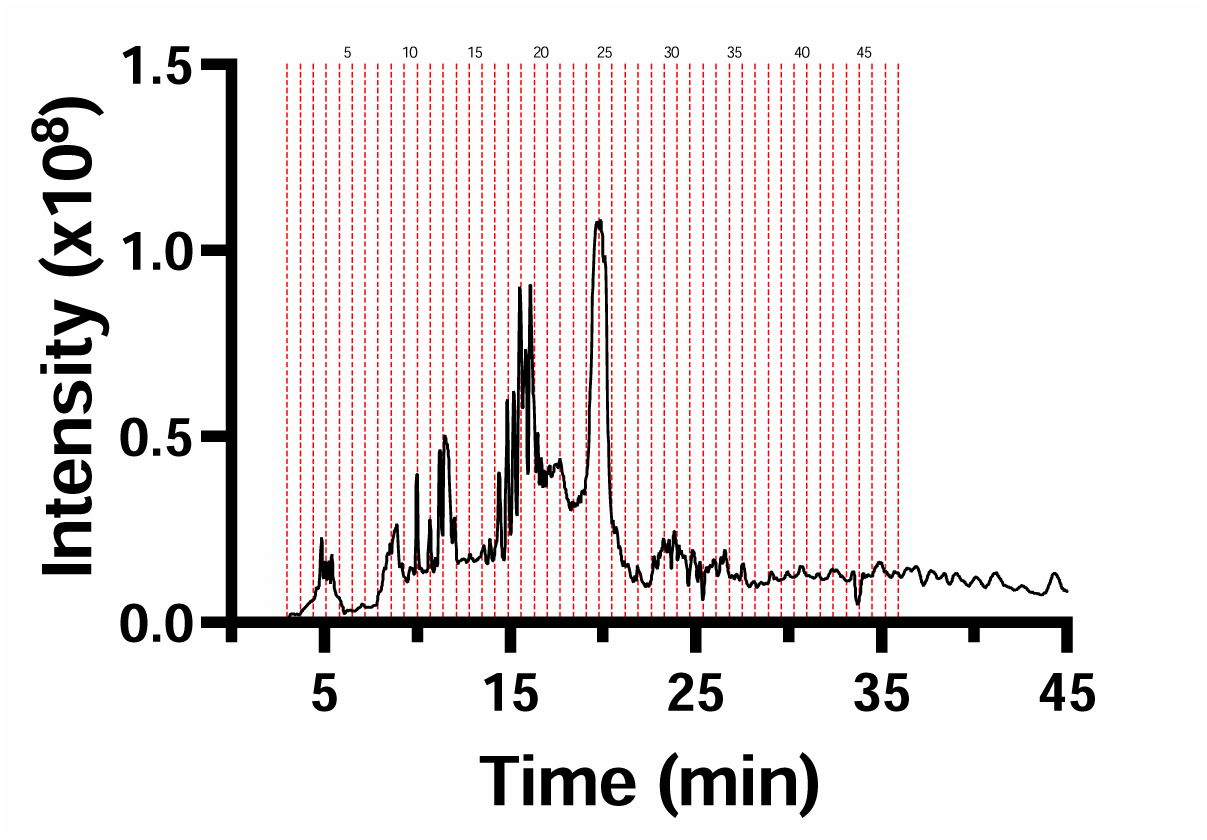
TIC of 900 μL injection of 2x pooled fecal matter by semi-preparative RPLC. Forty-seven fractions, corresponding to a 0.7-minute collection interval, were prepared for CSH and HILIC second-dimension separations.

**Figure 7.**
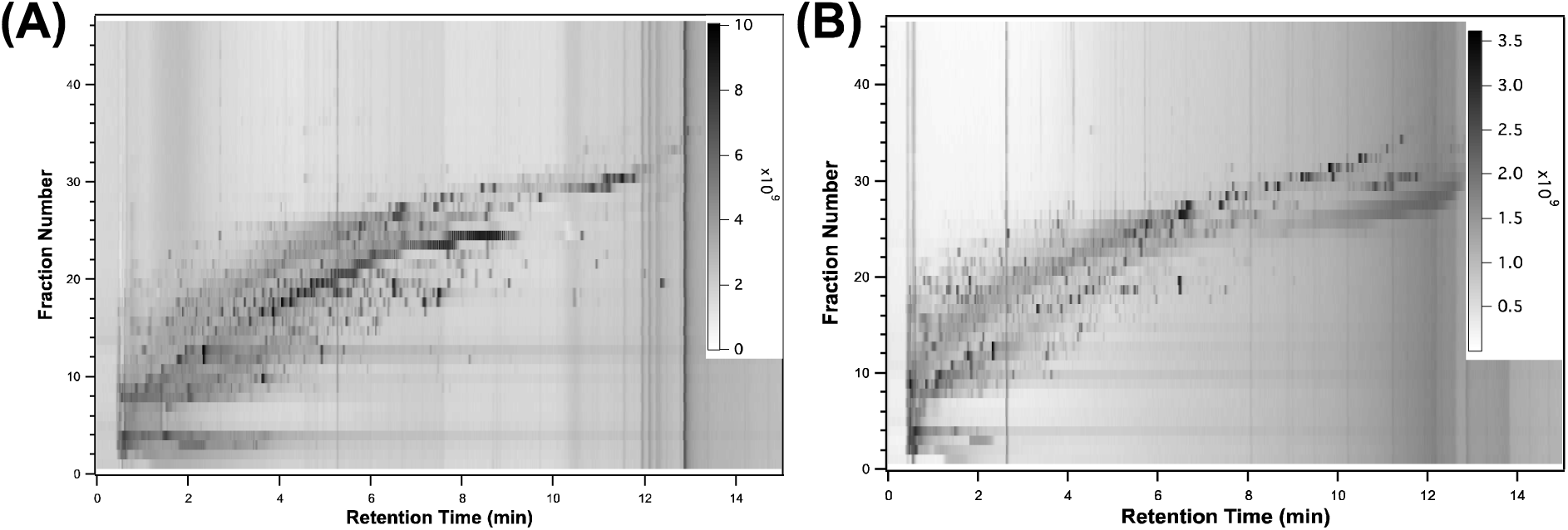
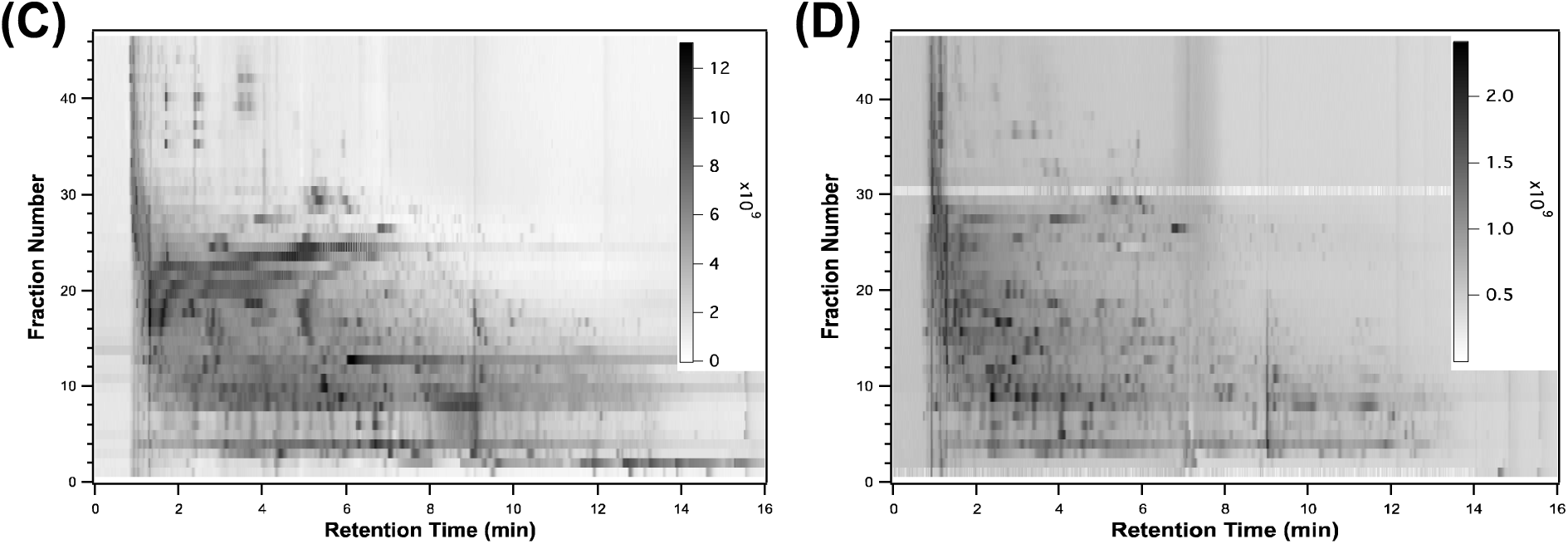
Two-dimensional TICs derived from LC-MS analysis of fractionated pooled fecal extract using (A) CSH positive mode, (B) CSH negative mode, (C) HILIC positive mode, and (D) HILIC negative mode methods. The logarithm of TIC intensity is represented by grayscale.

**Figure 8.**
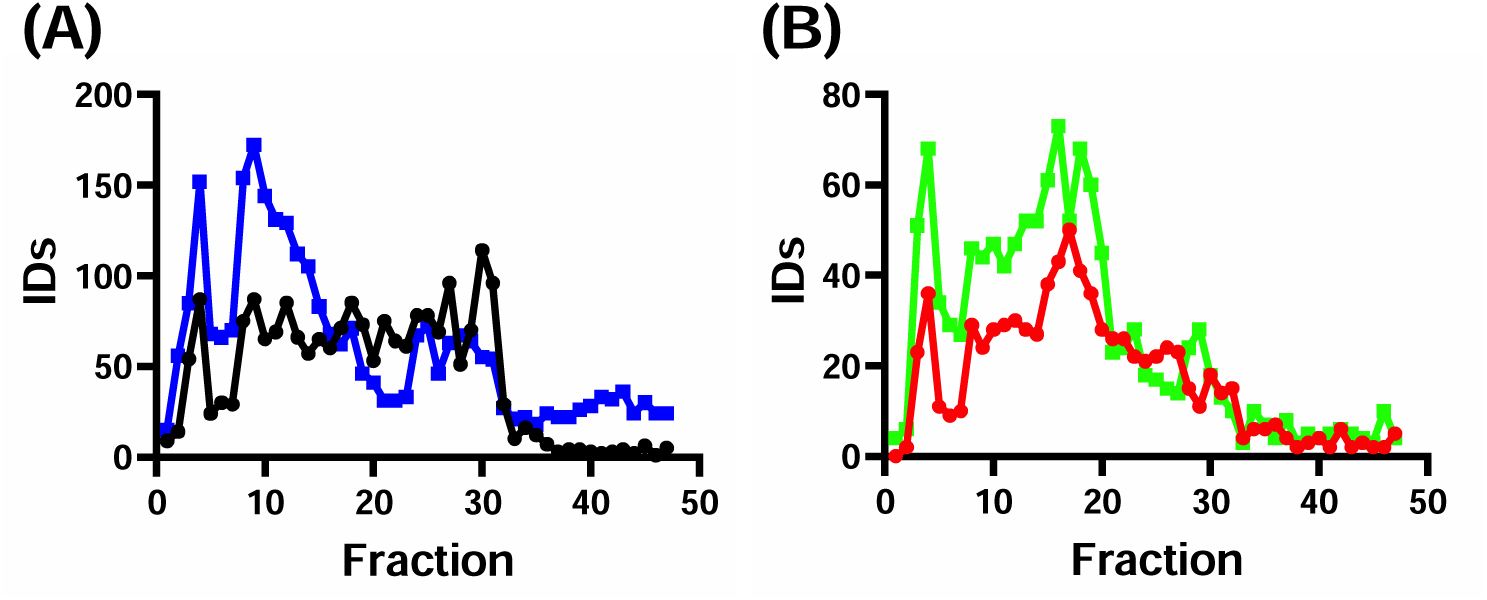
The number of compounds identified per fraction for (A) positive (black and blue) and (B) negative mode (red and green) with CSH and (circles) HILIC (squares) 2^nd^ dimension methods.

**Figure 9.**
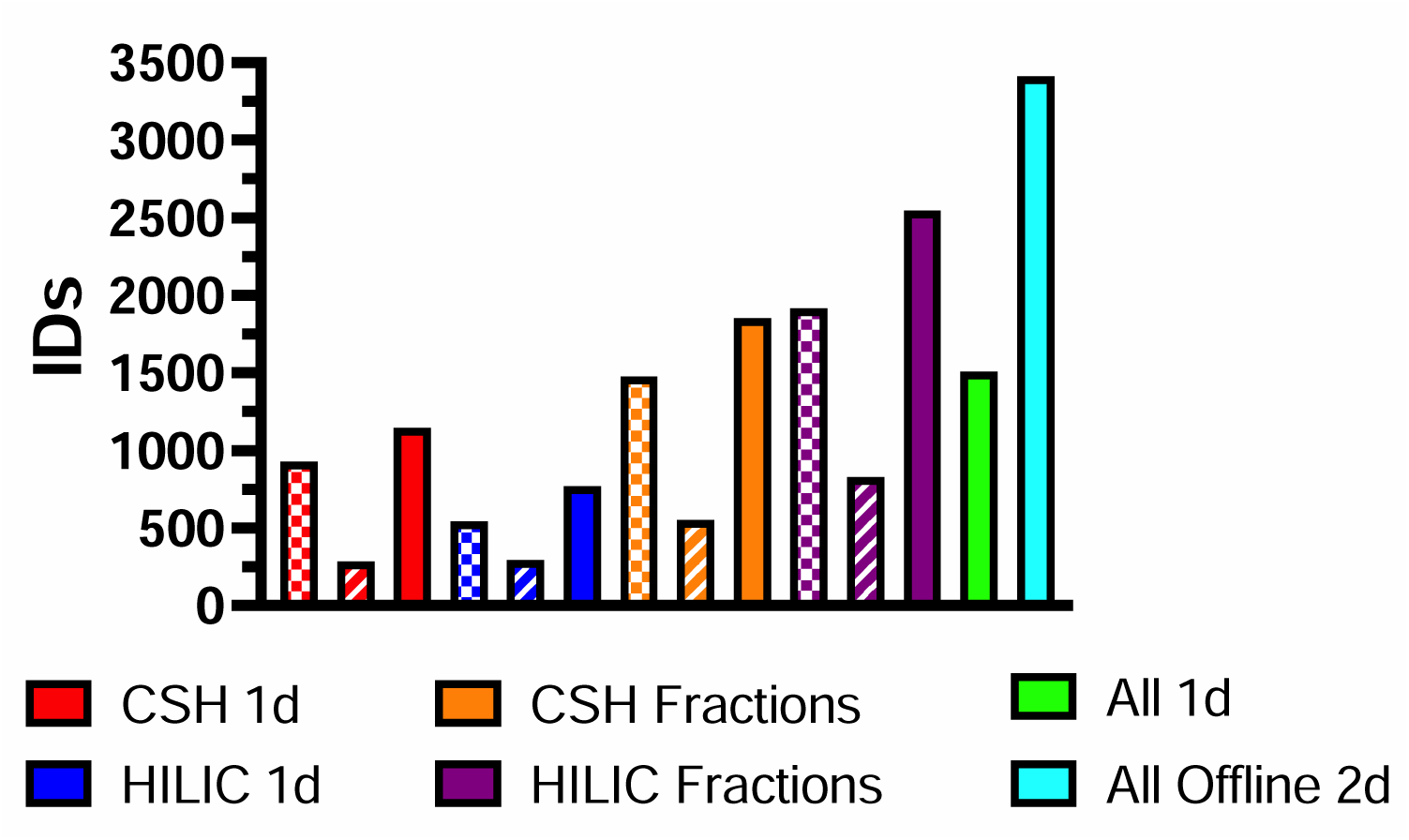
The total number of unique identified compounds observed by each method. Checkered and striped fill patterns are for positive and negative mode methods, respectively. Solid bars are a combination of both ion modes.

Compared to 1d separations, 2.25 times as many metabolites were identified by the offline 2d approaches, and a much higher number of MS/MS features were detected with a comparable proportion of unknowns (Figure 10). Compared to 1d methods, a larger proportion of the identifications achieved with the offline 2d methods were classified to MSI1 (11.5% to 10.8%) and MSI2A (59.7% to 46.8%) confidence levels. We have observed previously that the use of multi-run precursor ion exclusion as a strategy for increasing the number of compounds identified also tends to yield a higher total number of unidentified (MSI4) features, resulting in a higher proportion of unknowns even as the total number of identified compounds grew.^9^ In this study, however, the offline 2d methods identified a comparable proportion of the unknown MS/MS features as 1d methods, suggesting that the reduced sample complexity afforded by fractionation may help avoid this issue. Average precursor ion purity, a metric of how many different ions exist within the quadrupole isolation window of the MS scan, was lower (i.e., less “pure”) for 1D separations and had broader distributions for all identification levels (Figure 11). As precursor ion purity decreased, so did identification confidence; higher proportions of MSI4 features for 1D methods may also partly explain this observation.

**Figure 10.**
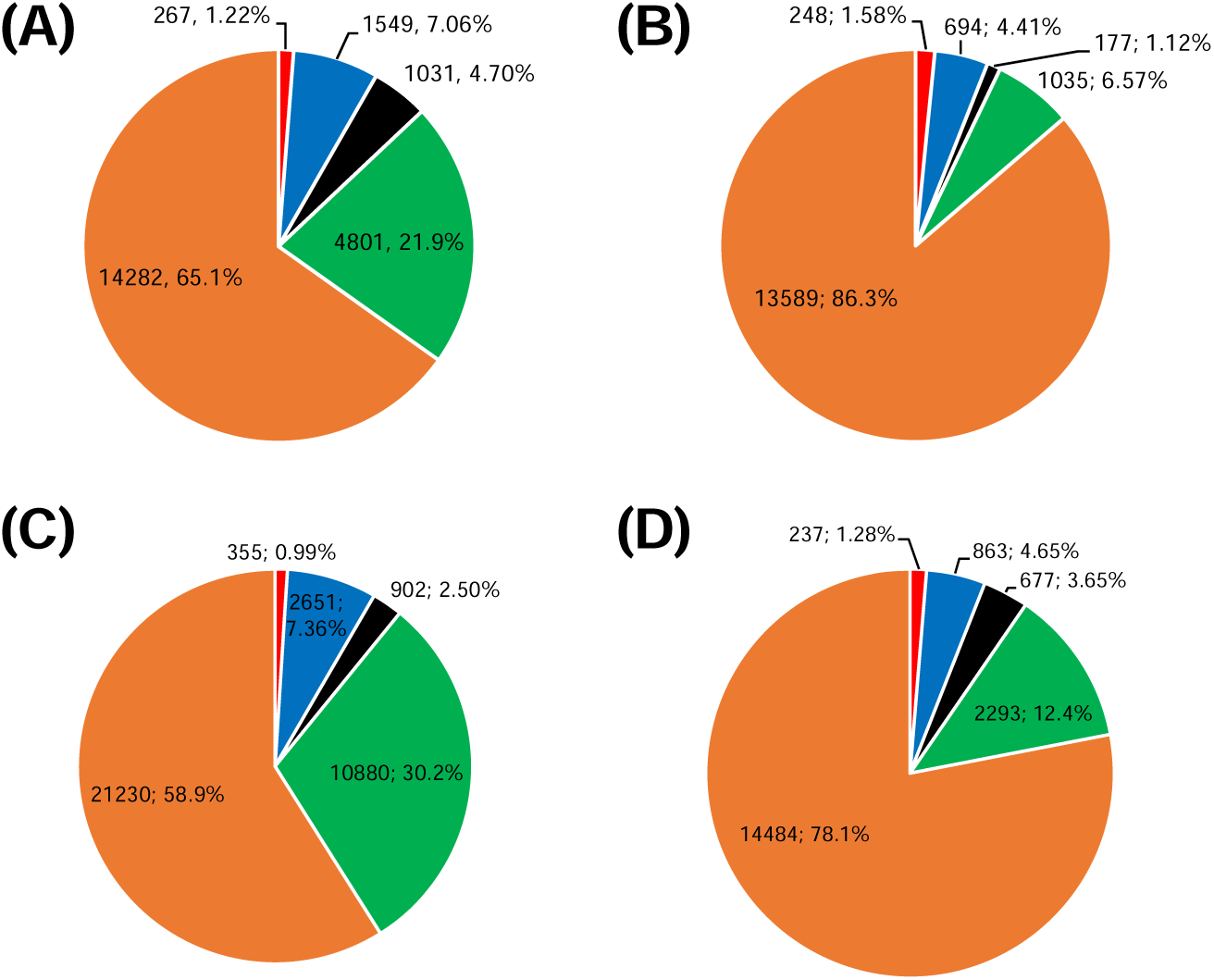
MSI level feature distribution for fractions collected via (A) CSH positive mode, (B) CSH negative mode, (C) HILIC positive mode, and (D) HILIC negative mode methods. The number of MS/MS features and overall percentage are displayed. ID levels MSI1, MSI2A, MSI2B, MSI3, and MSI4 are shown in red, blue, black, green, and orange.

**Figure 11.**
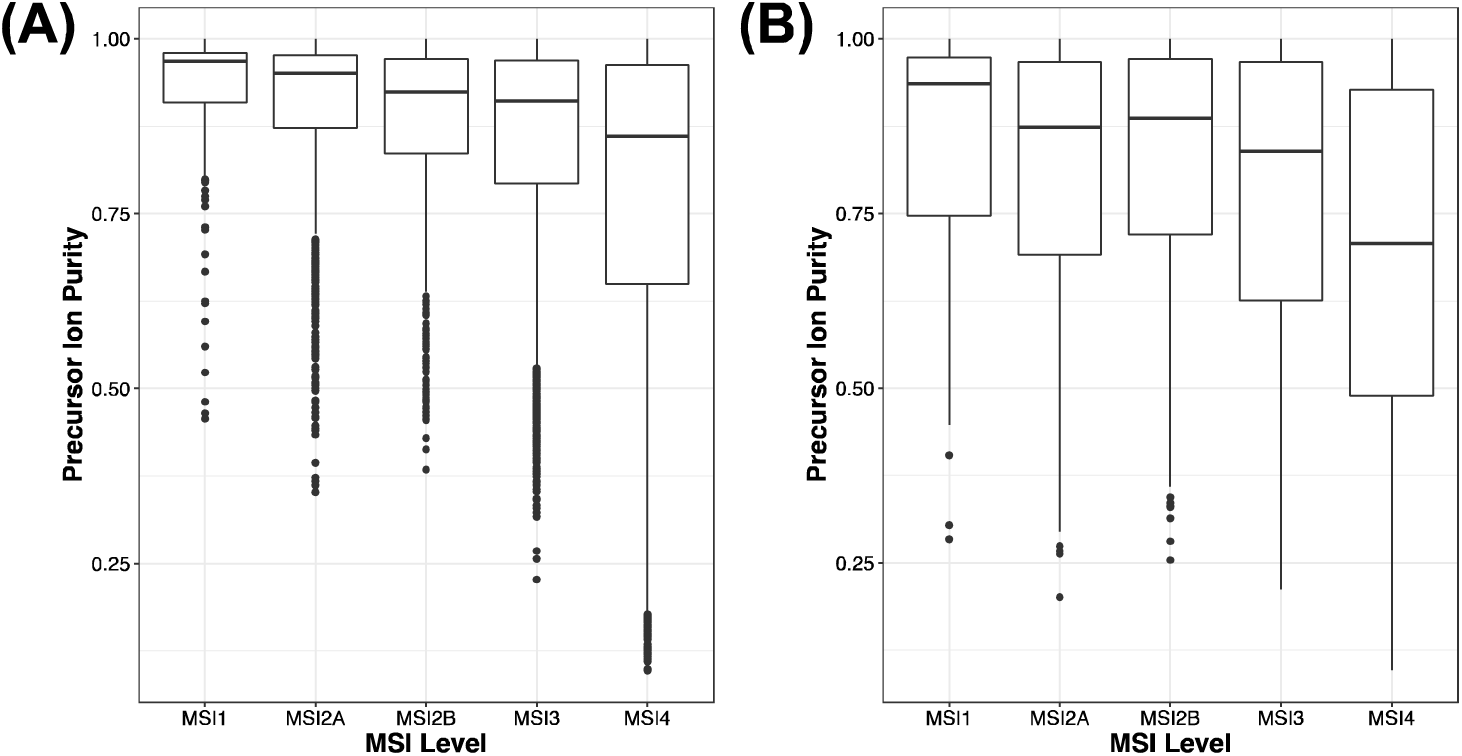
Precursor ion purity by MSI level for (A) fractions and (2) one-dimensional separation of a pooled fecal extract using the CSH positive mode method.

Iterative LC-MS/MS acquisition of the fractions was not performed to keep instrument run time under ∼16 hours for each separation and ionization mode combination. However, the potential benefit of using iterative MS/MS for deeper annotation of individual fractions was investigated using several fractions of varying abundance and regions of the first-dimension chromatogram (Figure 12). With each iterative injection, more metabolites were identified in each fraction, suggesting that further gains in compound identification can still be achieved if sample and run-time are not limited. Notably, even the first injection of an iterative MS/MS sequence resulted in the identification of more compounds than were observed in a typical single-injection, non-iterative data-dependent acquisition MS/MS worklist. This can be attributed to the creation of a dynamic exclusion list via injection of a blank sample before iterative sample injection sequences. In either case, more MS/MS spectra were acquired, fewer background ions were fragmented, and more metabolites were successfully identified using iterative acquisition.

**Figure 12.**
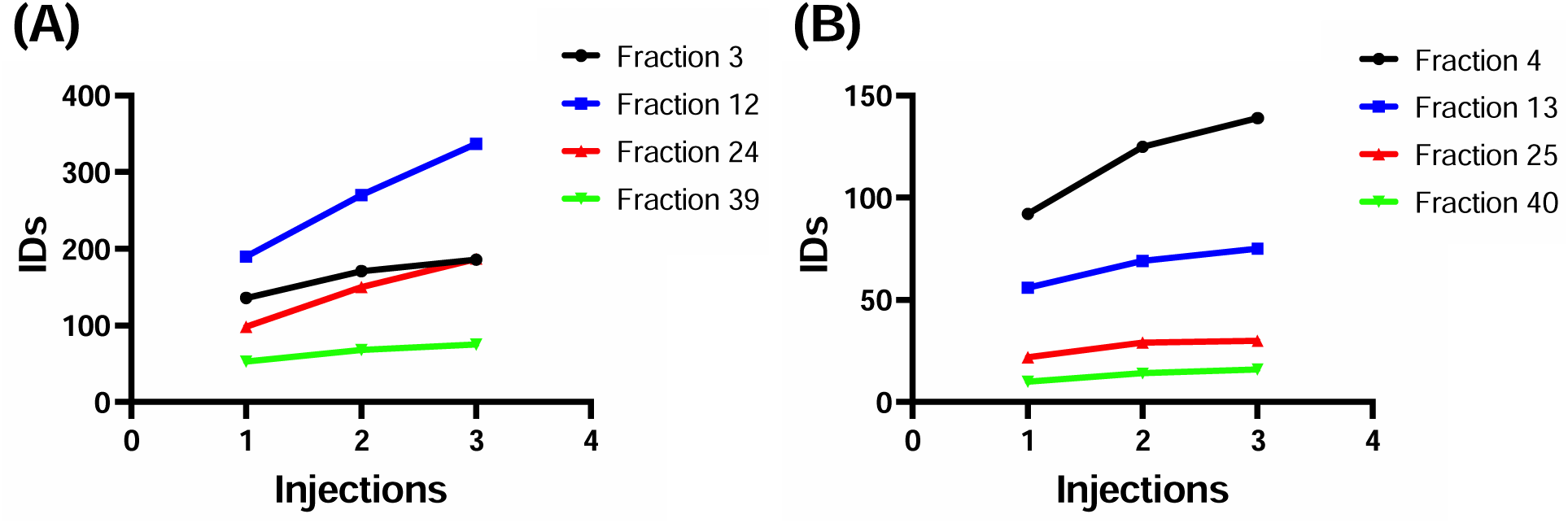
The number of cumulative unique identifications with three iterative injections for selected fractions by (A) HILIC positive mode and (B) HILIC negative mode second dimension methods.

Many metabolites were identified in multiple methods, including both 1d and 2d approaches, but 2261 identifications were unique to the offline 2d methods (Figure 13). Additional sample manipulation for 2d approaches (drying/reconstitution) and slightly different thresholds for removal of high-scoring database matches by iterative methods (creation of an exclusion list) and fractionation methods (metabolites in too many fractions) resulted in 360 metabolites that were only detected by 1d methods. Thus, to maximize total compound identification, the unfractionated sample should always be analyzed in addition to performing separate analyses of each fraction.

**Figure 13.**
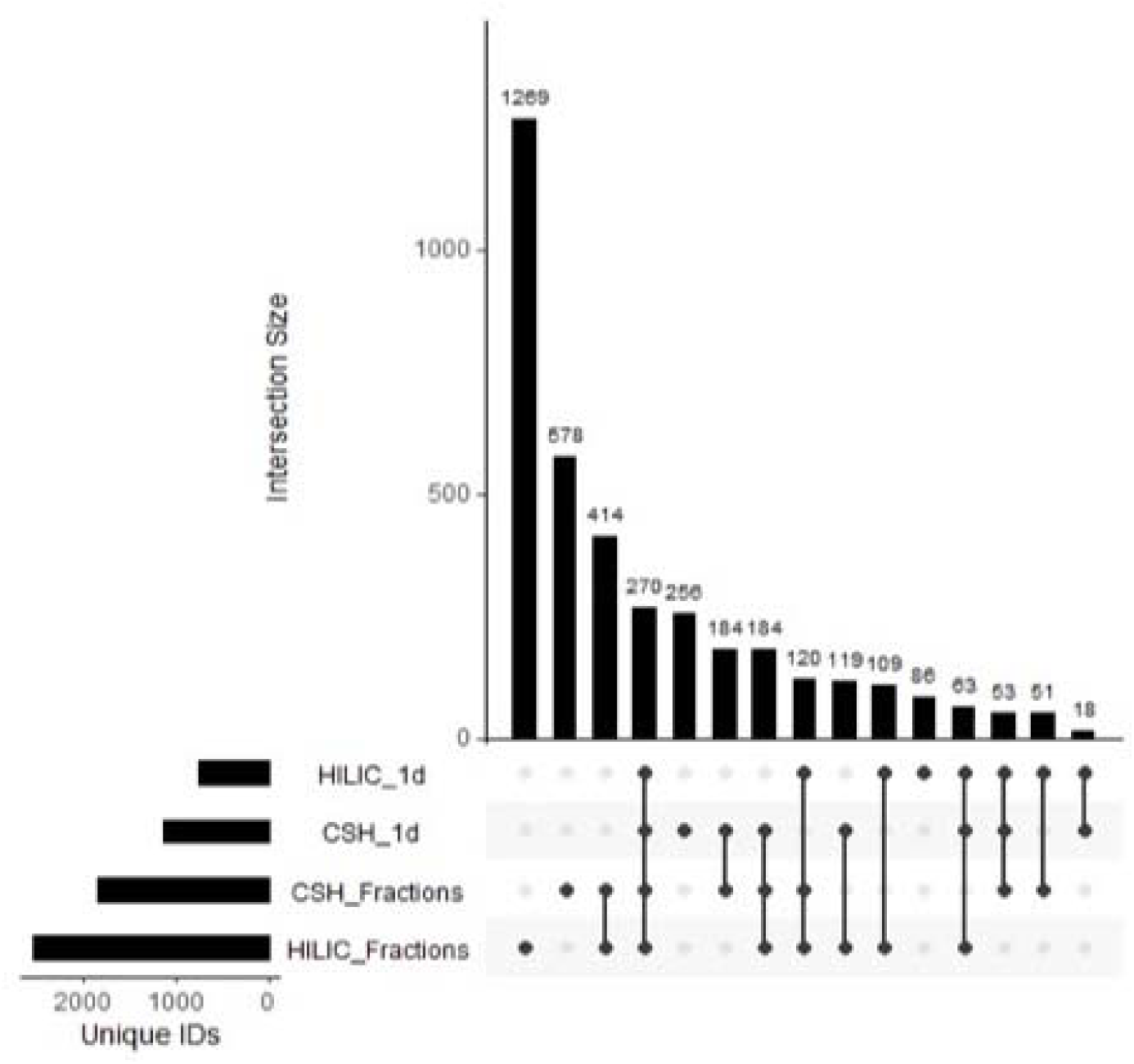
Overlap of identifications made by one-dimensional and two-dimensional methods. Identifications from positive and negative mode acquisition were combined before assessing overlap. Dots in the matrix signify the methods each group of identifications was found with.

### Evaluating Coverage of the Fecal Metabolome

Compound class information for all unique metabolite identifications and MSI3 annotations from a pooled fecal extract is shown in Figure 14. The most common chemical classes (superclass level for ClassyFire ontology) observed were lipid and lipid-like molecules (26.80%), organoheterocyclic compounds (20.11%), and organic acids and derivatives (17.28%). While lipid and lipid-like molecules was the most common superclass observed, it exhibited the lowest improvement from 1d methods in the number of unique identifications and compound class annotations achieved with offline 2d approaches (Figure 15). The greatest improvement in unique compound class annotations with fractionation and preconcentration was for benzenoids and nucleosides/nucleotides, which both demonstrated over a 3-fold increase from 1d methods. A substantial proportion of features in the dataset remained unidentified; while some represent degeneracies, adducts, and contaminants, another portion represents previously unannotated or unreported fecal metabolites. Strategies for the identification of such unknowns are discussed in more detail below.

**Figure 14.**
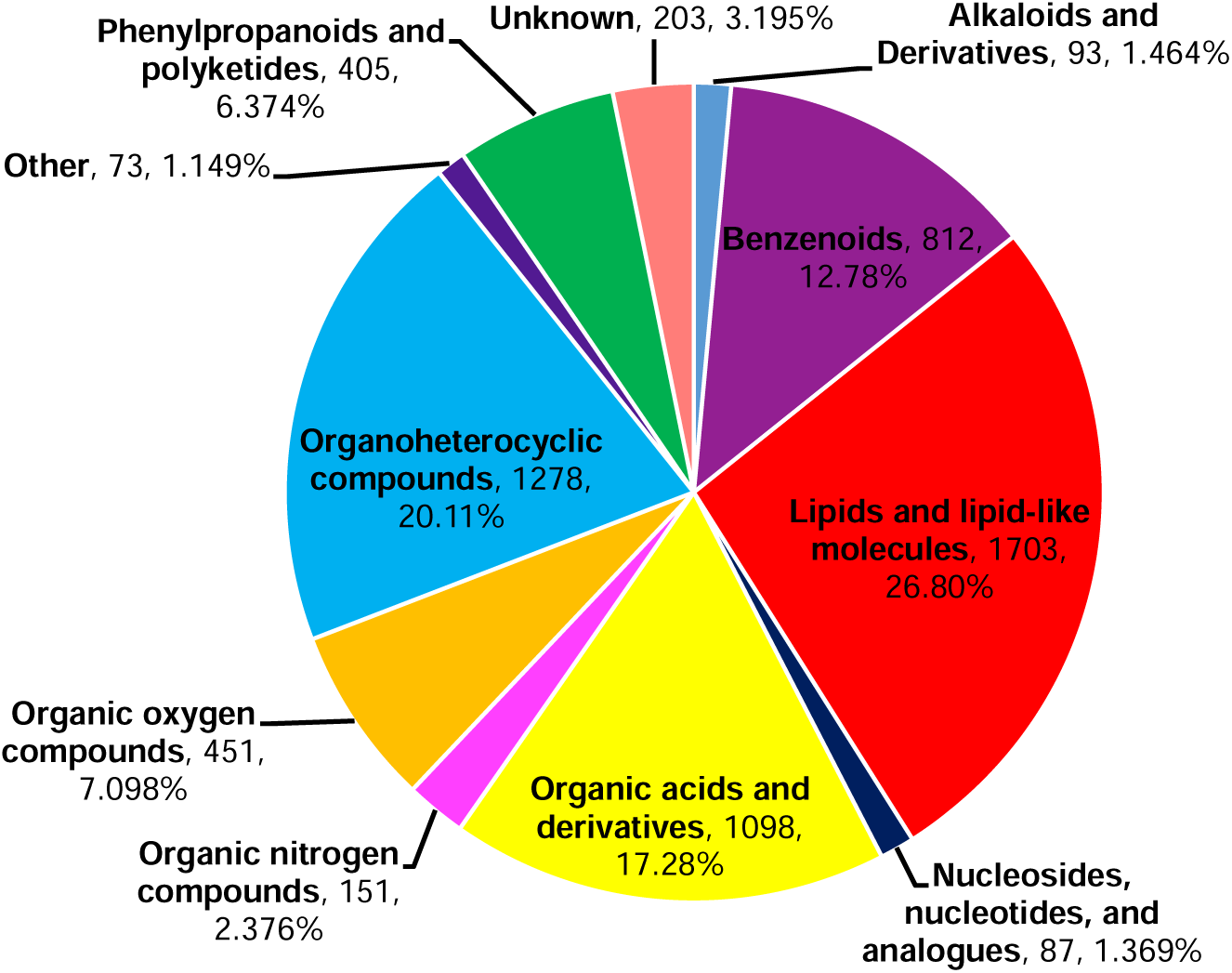
Compound class breakdown of all unique InChIKeys assigned an MSI1, MSI2A, MSI2B, and MSI3 identification level.

**Figure 15.**
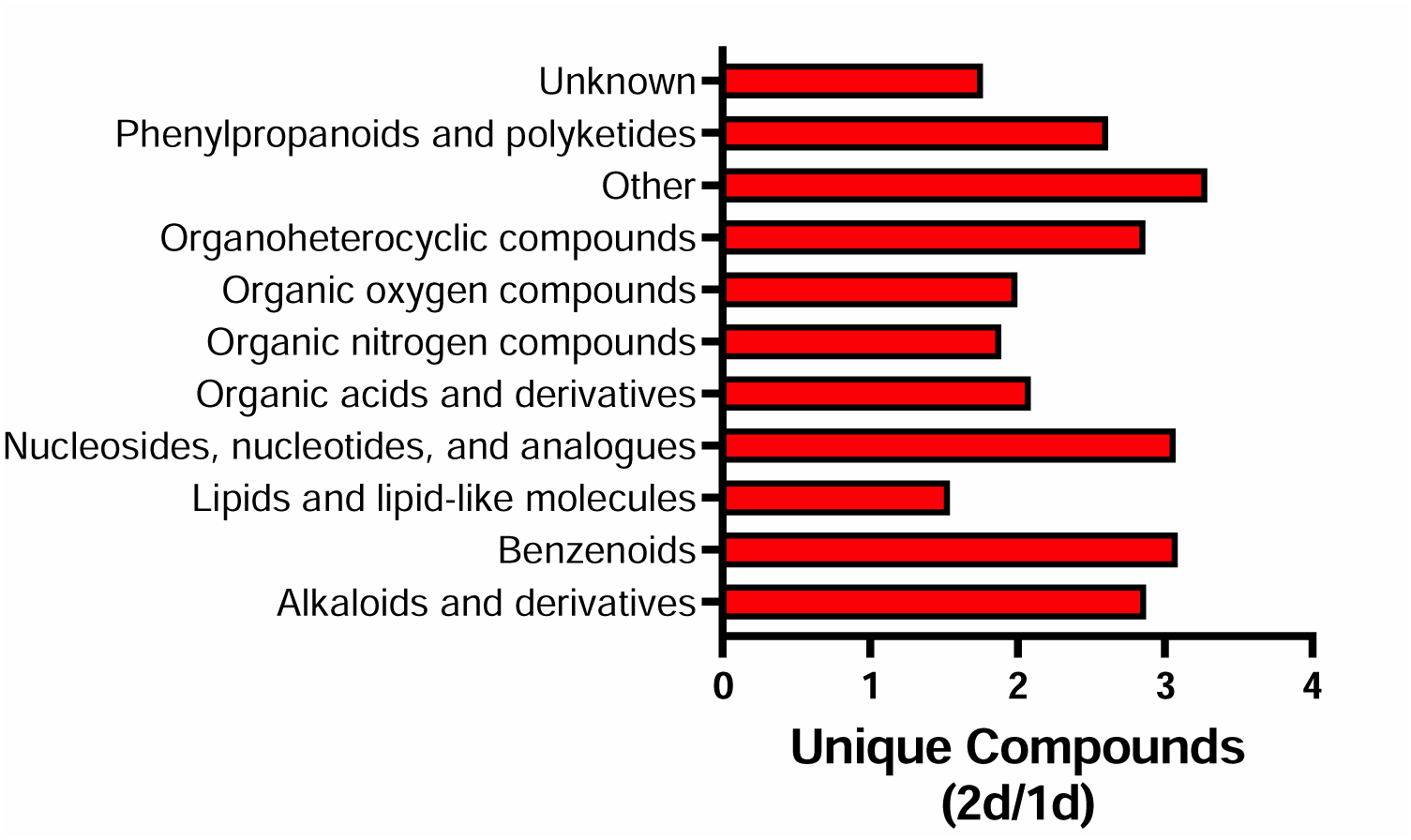
The ratio of all unique InChIKeys assigned an MSI1, MSI2A, MSI2B, and MSI3 identification level by compound class for offline 2d and 1d methods.

### Discovering New Metabolites

Fractionating and preconcentrating metabolites improved the signal and spectral quality of many unknown features. However, in spite of the improved chromatographic methods approaches utilized in this report, our data collection improvements only translated into enhanced identification performance if a library entry existed for the unknown metabolite. MS/MS spectral libraries continue to expand rapidly but remain incomplete. Continued improvements with *in silico* based structure prediction tools, including SIRIUS, CANOPUS, and COSMIC, may help prioritize and identify unknown features.^53–55^ When assessing our remaining unknown significant features with these tools, a high scoring (0.865 COSMIC score) proposed identification of a bile acid was achieved (Figure 16). MS/MS spectra for 7-oxoglycodeoxycholic acid did not exist in the databases searched. To date, 7-oxoglycodeoxycholic acid has never been reported as an identified metabolite in data uploaded to Metabolomics Workbench but is a plausible bacteria-produced secondary bile acid.^56^

**Figure 16.**
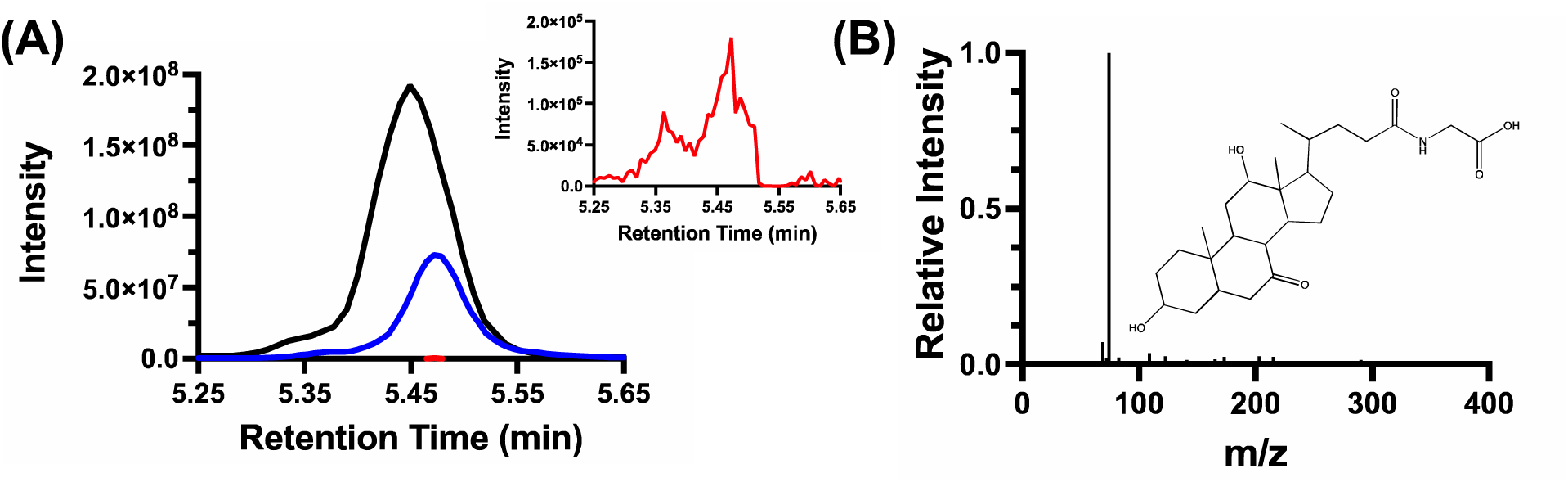
Significant feature from HILIC negative mode experiments. (A) extracted ion chromatograms of 426.2861 m/z from 1d separations of two pooled fecal samples (red, blue) compared to fraction 25 (black). (B) Experimental MS/MS and proposed identification 7-oxoglycodeoxycholic acid.

Elucidation of unknowns to annotate chemical classes such as bile acids could benefit from neutral loss searching strategies, as other conjugations than glycine and taurine may exist. For remaining unknown features that current *in silico* and data searching strategies yield no results, collecting nuclear magnetic resonance (NMR) data may be possible with our 2d approach. Low to mid mM concentrations are typically required for ^1^H NMR identification, but even effluent from conventional LC approaches has yielded promising results with lipids.^57^

## CONCLUSION

In this study, we demonstrated that by collecting and preconcentrating fractions from a semi-preparative first-dimension separation before an analytical second-dimension separation, the number of unique features identifiable in a pooled human feces extract by MS/MS spectral search increased 2.25-fold compared to several iterative 1d LC-MS/MS runs. Despite improvements in metabolite identification performance, many unknown features remain in our data. *In silico* and NMR-based approaches may provide complementary information helpful in annotating more unknown metabolites. Our study represents one of the most in-depth and careful characterizations of the fecal metabolome to date. Our strategy of using multidimensional separations to enhance compound identification performance in metabolomics should be equally applicable to other sample types. Current efforts are being directed toward evaluating its performance in the context of multiple human diseases and treatments.

## AUTHOR INFORMATION

### Author Contributions

B.G.A, C.M.T., R.T.K., and C.R.E. designed the study. Data processing and compound annotation were performed by B.G.A. R.H. prepared and annotated retention times of the MetaSci standards library. A.R. developed *MetIDTracker* software used for database searching and data alignment. Fecal matter patient samples were collected by M.K.D., S.K.M., and A.G. Sample preparation and the collection of LC-MS experimental data were performed by B.G.A. and C.R.E. All authors contributed to the writing and editing of the manuscript.

## ACKNOWLEDGEMENTS

This work was supported by the Common Fund Metabolomics program, grant U2CES030164 (C.R.E. and Alexey Nesvizhskii), grant P41LGM108538 (Joshua Coon), and grant R01DK101473-01A1 (R.T.K.), all of which are funded by the National Institutes of Health (NIH).

## Notes

### Competing Interest Statement

The authors have declared no competing interest.

